# Cold exposure distinctively modulates parathyroid and thyroid hormones in cold-acclimatized and non-acclimatized humans

**DOI:** 10.1101/2020.01.16.906081

**Authors:** Zuzana Kovaničová, Tímea Kurdiová, Miroslav Baláž, Patrik Štefanička, Lukáš Varga, Oana C. Kulterer, Matthias J. Betz, Alexander R. Haug, Irene A. Burger, Florian W. Kiefer, Christian Wolfrum, Barbara Ukropcová, Jozef Ukropec

**Affiliations:** Institute of Experimental Endocrinology, Biomedical Research Center, Slovak Academy of Sciences, Bratislava, Slovakia; Institute of Pathophysiology, Faculty of Medicine, Comenius University, Bratislava, Slovakia; Department of Health Sciences and Technology, Institute of Food, Nutrition and Health ETH Zürich, Zürich, Switzerland; Department of Otorhinolaryngology – Head and Neck Surgery, Faculty of Medicine Comenius University and University Hospital Bratislava, Bratislava, Slovakia; Division of Endocrinology and Metabolism, Department of Medicine III, Medical University of Vienna, Vienna, Austria; Department of Endocrinology, Diabetes & Metabolism, University Hospital of Basel, Basel, Switzerland; Department of Biomedical Imaging and Image-guided Therapy, Division of Nuclear Medicine, Medical University of Vienna, Vienna, Austria; Department of Nuclear Medicine, University Hospital Zürich, Zürich, Switzerland; Faculty of Physical Education and Sports, Comenius University, Bratislava, Slovakia

**Keywords:** parathyroid & thyroid hormones, cold exposure, non-shivering thermogenesis, brown adipose tissue, 18F-FDG-PET, cold acclimatization

## Abstract

**Context:** Cold-induced activation of thermogenesis modulates energy metabolism, but the role of humoral mediators is not completely understood.

**Objective:** To investigate the role of parathyroid and thyroid hormones in acute and adaptive response to cold in humans.

**Design:** Cross-sectional study examining acute response to ice-water swimming and to experimental non-shivering thermogenesis (NST) induction in individuals acclimatized and non-acclimatized to cold. Seasonal variation in energy metabolism of ice-water swimmers and associations between circulating PTH and molecular components of thermogenic program in brown adipose tissue (BAT) of neck-surgery patients were evaluated.

**Setting:** Clinical Research Center.

**Patients, Participants:** Ice-water swimmers (winter swim n=15, NST-induction n=6), non-acclimatized volunteers (NST-induction, n=11, elective neck surgery n = 36).

**Main Outcomes and Results:** In ice-water swimmers, PTH and TSH increased in response to 15min winter swim, while activation of NST failed to regulate PTH and lowered TSH. In non-acclimatized men, NST-induction decreased PTH and TSH. Positive correlation between systemic levels of PTH and whole-body metabolic preference for lipids as well as BAT 18F-FDG uptake was found across the two populations. Moreover, NST-cooling protocol-induced changes in metabolic preference for lipids correlated positively with changes in PTH. Finally, variability in circulating PTH correlated positively with *UCP1*/UCP1, *PPARGC1A* and *DIO2* in BAT from neck surgery patients.

**Conclusions:** Regulation of PTH and thyroid hormones during cold exposure in humans depends on the cold acclimatization level and/or cold stimulus intensity. Role of PTH in NST is substantiated by its positive relationships with whole-body metabolic preference for lipids, BAT volume and UCP1 content.

## Introduction

During evolution, humans have developed various cold-coping mechanisms, including shivering and non-shivering thermogenesis (NST) in skeletal muscle and/or brown adipose tissue (BAT) ^1^, with the principal thermogenic machinery in BAT represented by mitochondrial uncoupling protein 1 (UCP1). Importantly, human BAT could have similar thermogenic functionality to rodents relative to mitochondrial content ^2^ and several studies have observed metabolic effects associated with cold-induced BAT activation in humans, including increased energy expenditure, insulin sensitivity, lipolysis and fatty acid oxidation ^2–5^. Cold-induced metabolic activation of thermogenic processes could therefore potentially help control the energy imbalance and pathophysiological consequences of the nutrient overload present in obesity and metabolic disease ^6^.

Adipose organ plasticity allows for substantial morphological and functional transformation in adaptive response to repeated cold exposure or regular physical activity. This includes adipose tissue “browning” - the formation of multilocular adipocytes within white adipose tissue (WAT), indicating enhanced capacity for energy dissipation ^7^. It has been demonstrated that repeated cold exposure can induce BAT activity in lean adults ^8^, healthy individuals with obesity ^9^ or patients with type 2 diabetes ^10^. Therefore, regular cold exposure could represent an efficient means of the adipose tissue thermogenic program activation in most individuals, independently of age or adiposity, which are usually negatively associated with the amount of BAT ^11–12^.

We studied systemic humoral mediators of thermogenesis in humans well acclimatized to cold. We hypothesized that this specific population is more likely to possess efficient thermogenic mechanisms, including thermogenically active BAT, and that systemic mediators of the thermogenic process could be altered following acute exposure to cold. We therefore explored the effects of acute and seasonal severe cold exposure on circulating hormones and metabolites that might be involved in the development of cold-coping mechanisms that enable efficient acute and chronic cold acclimation in individuals that regularly engage in outdoor ice-cold water swimming (<5°C). We focused (i) on thyroid hormone axis due to its role in energy homeostasis and its potential to induce adipose tissue browning in both animals and humans ^13–14^ and (ii) on parathyroid hormone (PTH) that has been recently shown to induce adipose tissue browning in rodents and in cultured human adipocytes ^15–16^. We also investigated seasonal differences in baseline (unstimulated) levels of key molecules regulating or reflecting metabolic preference, insulin sensitivity and thermogenic response to ice-water swimming together with anthropometric and biochemical characteristics (Cohort 1).

Next, we compared effects of ice-cold water swimming (cold stress) and controlled mild cold exposure (NST activation) in cold acclimatized individuals (Cohort 2) on systemic levels of PTH and thyroid hormones. We explored the relationships of PTH with BAT volume and glucose uptake, whole-body energy metabolism and metabolic substrate preference. In parallel we also studied effects of NST activation in healthy young lean non-acclimatized men (Cohort 3). The relationships of PTH and thyroid hormones with molecular hallmarks of BAT thermogenic potential, i.e. UCP1 mRNA and protein, was explored in deep-neck BAT samples from patients undergoing elective neck surgery, who were neither acclimatized nor acutely exposed to cold (Cohort 4).

## Methods

### Study population and protocol

All participants were fully informed about the study protocol and signed a written informed consent prior entering the study. The study was approved by the Ethics Committee of the Faculty of Medicine Comenius University & University Hospital Bratislava (Cohort 1 and 3), by the ethics committees of the Medical University of Vienna (Cohort 2) and the canton of Zürich (Cohort 3). The Study conforms to Declaration of Helsinki, as amended in 2013, and with the standards of the International Conference on Harmonization (ICH) & Good Clinical Practice (GCP).

#### Cohort 1

We recruited 15 middle-aged ice-water swimmers who had engaged in regular outdoor cold-water swimming for at least past 6 months (Table 1). At the time of the study, all participants lived in continental European climate and were adequately trained and acclimatized to endure ≥10min of swimming in <5°C cold water. The winter ice-water swimming event, organized by a local ice-water swimming club (http://www.ladovemedvede.sk/), began with approximately 30min acclimation to outdoor temperature (0°C) in light clothing (swimsuit, T-shirt), followed by swimming in 2.6°C water (15min on average) in the Danube river in Bratislava, Slovakia (February), wearing only a swimsuit (no thermal protection) and a head cover. Blood was collected indoors (25°C) from a cubital vein approximately 1h before and 10-30min after swimming in the river. Food intake prior to the event was not restricted, but participants were asked to refrain from food consumption before the blood collection after swimming. In the following month (average daily air temperature of sampling period: 7.2°C), volunteers visited our clinical unit after an overnight fast for additional blood sampling and assessment of body composition (bioelectrical impedance, Omron BF511, Japan), blood pressure and pulse (Omron 907, Japan) and cold-hardening habits (questionnaire). Seven months later, at the end of summer (average daily air temperature of sampling period: 18.3°C), phenotyping and blood collection were repeated (Table 1). Design of the study is detailed in Figure 1. Self-reported weight and height values from one volunteer who did not attend clinical phenotyping were used in the analyses. One participant was treated for hypertension (monotherapy with beta-blockers).

**Table 1:**
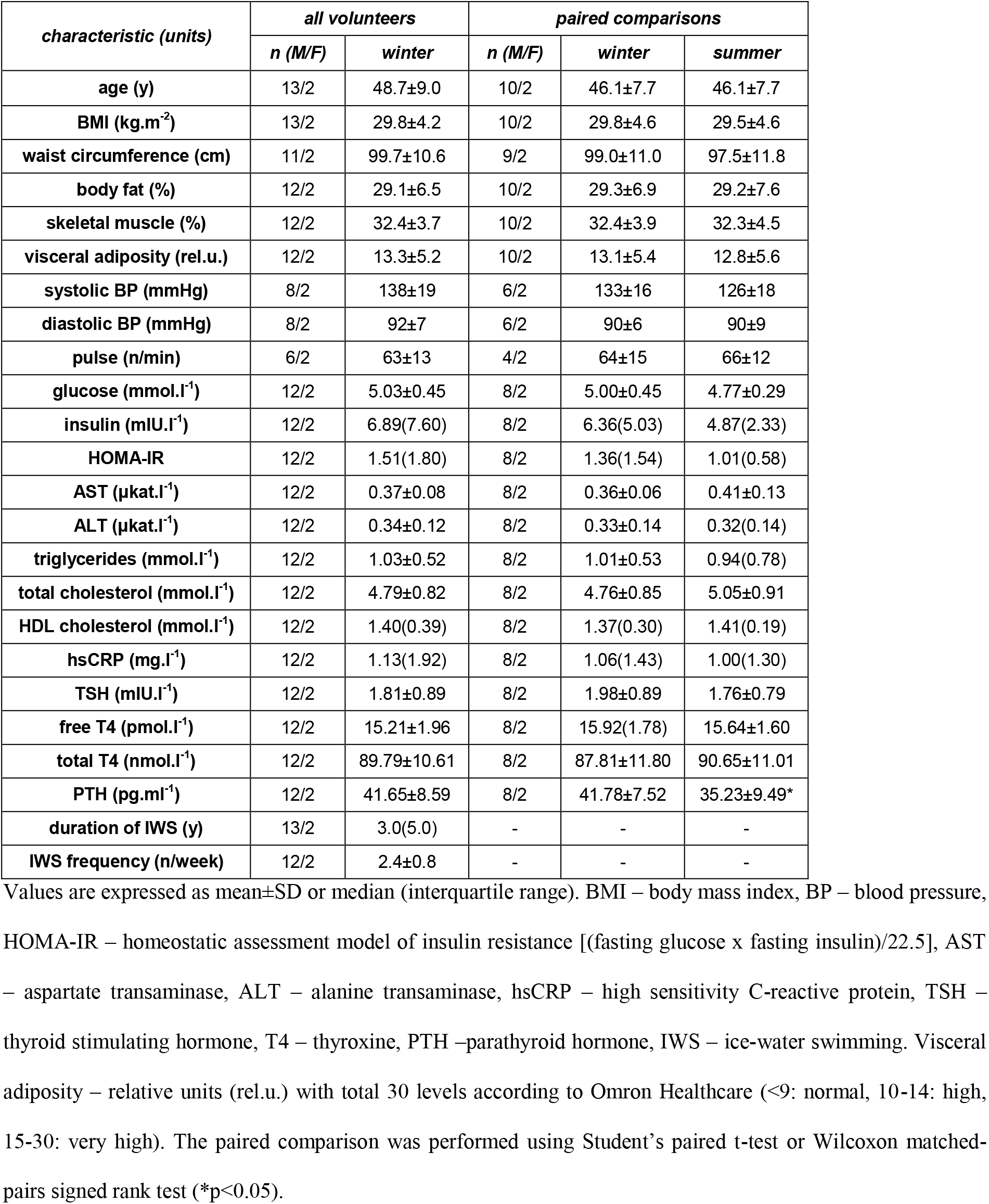
Characteristics of Cohort 1 in winter (all) and seasonal paired comparison on a subset of volunteers who underwent clinical examination both in winter (February) and late summer (September) period.

| characteristic (units) | all volunteers |  | paired comparisons |  |  |
| --- | --- | --- | --- | --- | --- |
|  | n (M/F) | winter | n (M/F) | winter | summer |
| age (y) | 13/2 | 48.7±9.0 | 10/2 | 46.1±7.7 | 46.1±7.7 |
| BMI (kg.m<sup>-2</sup>) | 13/2 | 29.8±4.2 | 10/2 | 29.8±4.6 | 29.5±4.6 |
| waist circumference (cm) | 11/2 | 99.7±10.6 | 9/2 | 99.0±11.0 | 97.5±11.8 |
| body fat (%) | 12/2 | 29.1±6.5 | 10/2 | 29.3±6.9 | 29.2±7.6 |
| skeletal muscle (%) | 12/2 | 32.4±3.7 | 10/2 | 32.4±3.9 | 32.3±4.5 |
| visceral adiposity (rel.u.) | 12/2 | 13.3±5.2 | 10/2 | 13.1±5.4 | 12.8±5.6 |
| systolic BP (mmHg) | 8/2 | 138±19 | 6/2 | 133±16 | 126±18 |
| diastolic BP (mmHg) | 8/2 | 92±7 | 6/2 | 90±6 | 90±9 |
| pulse (n/min) | 6/2 | 63±13 | 4/2 | 64±15 | 66±12 |
| glucose (mmol.l<sup>-1</sup>) | 12/2 | 5.03±0.45 | 8/2 | 5.00±0.45 | 4.77±0.29 |
| insulin (mIU.l<sup>-1</sup>) | 12/2 | 6.89(7.60) | 8/2 | 6.36(5.03) | 4.87(2.33) |
| HOMA-IR | 12/2 | 1.51(1.80) | 8/2 | 1.36(1.54) | 1.01(0.58) |
| AST (μkat.l<sup>-1</sup>) | 12/2 | 0.37±0.08 | 8/2 | 0.36±0.06 | 0.41±0.13 |
| ALT (μkat.l<sup>-1</sup>) | 12/2 | 0.34±0.12 | 8/2 | 0.33±0.14 | 0.32(0.14) |
| triglycerides (mmol.l<sup>-1</sup>) | 12/2 | 1.03±0.52 | 8/2 | 1.01±0.53 | 0.94(0.78) |
| total cholesterol (mmol.l<sup>-1</sup>) | 12/2 | 4.79±0.82 | 8/2 | 4.76±0.85 | 5.05±0.91 |
| HDL cholesterol (mmol.l<sup>-1</sup>) | 12/2 | 1.40(0.39) | 8/2 | 1.37(0.30) | 1.41(0.19) |
| hsCRP (mg.l<sup>-1</sup>) | 12/2 | 1.13(1.92) | 8/2 | 1.06(1.43) | 1.00(1.30) |
| TSH (mIU.l<sup>-1</sup>) | 12/2 | 1.81±0.89 | 8/2 | 1.98±0.89 | 1.76±0.79 |
| free T4 (pmol.l<sup>-1</sup>) | 12/2 | 15.21±1.96 | 8/2 | 15.92(1.78) | 15.64±1.60 |
| total T4 (nmol.l<sup>-1</sup>) | 12/2 | 89.79±10.61 | 8/2 | 87.81±11.80 | 90.65±11.01 |
| PTH (pg.ml<sup>-1</sup>) | 12/2 | 41.65±8.59 | 8/2 | 41.78±7.52 | 35.23±9.49* |
| duration of IWS (y) | 13/2 | 3.0(5.0) | - | - | - |
| IWS frequency (n/week) | 12/2 | 2.4±0.8 | - | - | - |

Values are expressed as mean±SD or median (interquartile range). BMI – body mass index, BP – blood pressure, HOMA-IR – homeostatic assessment model of insulin resistance [(fasting glucose x fasting insulin)/22.5], AST – aspartate transaminase, ALT – alanine transaminase, hsCRP – high sensitivity C-reactive protein, TSH – thyroid stimulating hormone, T4 – thyroxine, PTH –parathyroid hormone, IWS – ice-water swimming. Visceral adiposity – relative units (rel.u.) with total 30 levels according to Omron Healthcare (<9: normal, 10-14: high, 15-30: very high). The paired comparison was performed using Student's paired t-test or Wilcoxon matched-pairs signed rank test (\*p<0.05).

**Figure 1:**
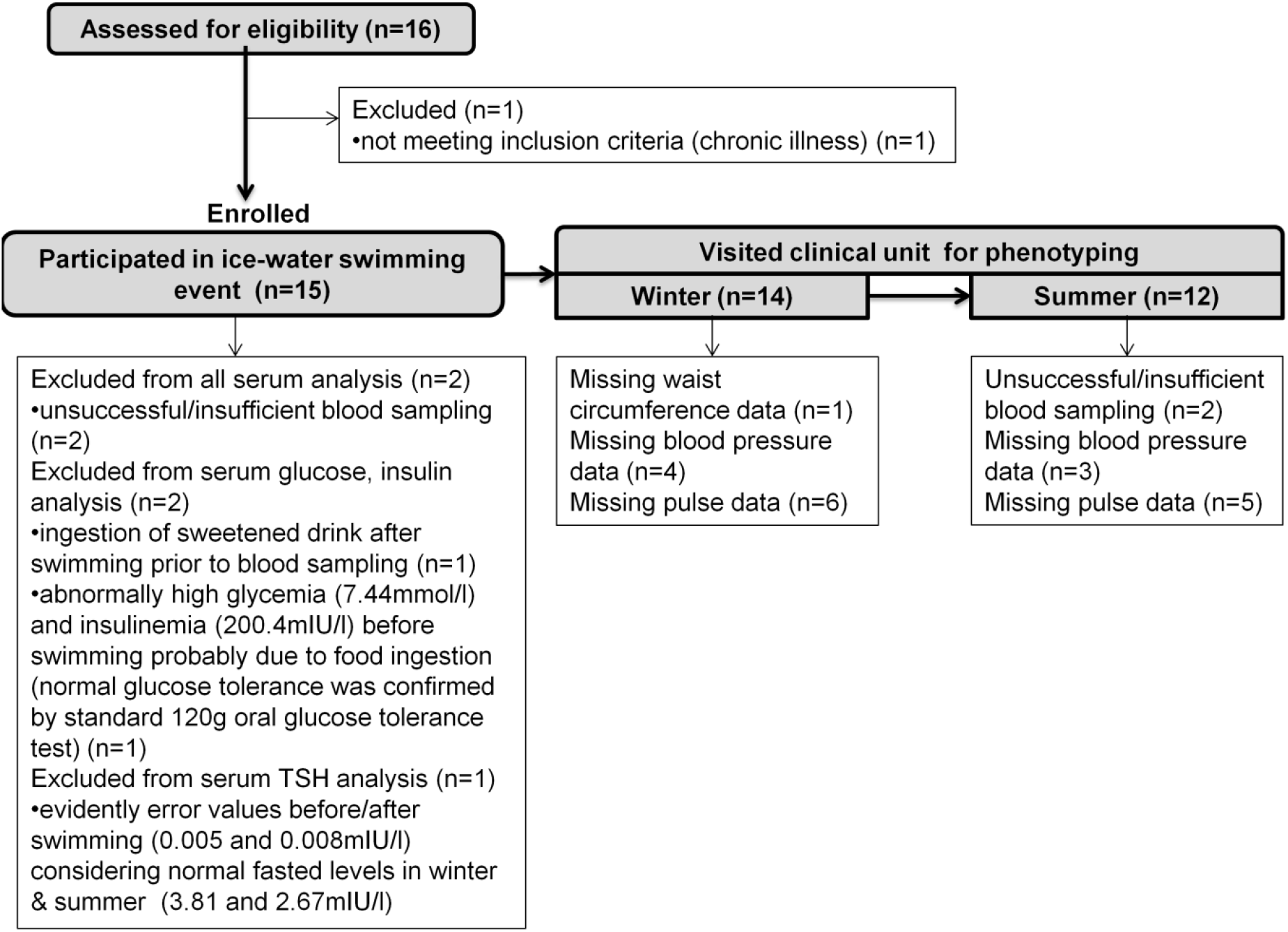
Study design for Cohort 1.

#### Cohort 2

This study was performed at the Medical University of Vienna. We used plasma samples and measurements from 6 participants (characteristics - Table 2) of the clinical study NCT02381483, who were all members of the ice-water swimming club (http://www.ladovemedvede.sk/) and two of them participated in Cohort 1. Study participants were examined at two separate study visits in the morning after an overnight fast (>10h). At study visit one, a blood draw was performed followed by an indirect calorimetry using a computerized open circuit indirect calorimetry (Quark RMR, Cosmed, Rome, Italy) in order to determine the resting energy expenditure at thermoneutrality (room temperature). Afterwards subjects received 2,5 MBq/kg^−1^ BW [^18^F]-FDG (max 350 MBq) and underwent the first [^18^F]-FDG PET/CT scan at room temperature (24°C) to detect any basal BAT activity using a Siemens Biograph 64 True Point PET/CT scanner (Siemens Healthcare Sector, Erlangen, Germany).The PET and CT images were acquired from the base of the skull until mid-thigh. On study day two a blood draw was performed, followed by an indirect calorimetry at thermoneutrality (room temperature). Then, a personalized cooling protocol was applied for a total of 150 min using a water-perfused cooling vest (CoolShirt Systems, Stockbridge, Georgia, USA). The water temperature was kept slightly above the shivering temperature (6,5 ± 3,4°C) and muscle activity was monitored by electromyography (OT Bioelettronica, Torino, Italy). After 90 min of cold exposure a second indirect calorimetry was performed followed by the administration of 2,5 MBq/kg^−1^ BW [^18^F]-FDG (max 350 MBq) and cold exposure continued for another 60 min until the PET/CT scan followed by blood sampling.

**Table 2:** The antropometric, metabolic and hormonal parameters of Cohorts 2 and 3 at warm (baseline) and non-shivering cold condition.

| *study group* | *cold-acclimatized (Cohort 2)* |  | *non-acclimatized (Cohort 3)* |  |
| --- | --- | --- | --- | --- |
| *n* | *5 males/1 female* |  | *11 males* |  |
| *condition* | *warm* | *cold* | *warm* | *cold* |
| age (years) | 37.2±4.4 | - | 22.7±2.6 | - |
| BMI (kg.m<sup>-2</sup>) | 25.1±5.0 | - | 23.4±1.9 | - |
| fat mass (kg) | 20.3±11.3 | - | 16.5±3.5 | - |
| lean body mass (kg) | 60.7±10.8 | - | 60.5±2.6 | - |
| RMR (kcal/day) | 1719.5±299.5 | 1830.3±293.7* | 2166.8±277.2 | 2424.6±320.8** |
| RQ (VCO<sub>2</sub>/VO<sub>2</sub>) | 0.81±0.04 | 0.77±0.06* | 0.78±0.08 | 0.84±0.07** |
| CIT (%) | - | 6.7±4.7 | - | 12.2±9.8 |
| BAT volume (cm<sup>2</sup>) | - | 137.6±118.2 | - | 56.8±37.8 |
| SUV mean/LBM (a.u.) | - | 2.9±0.54 | - | 4.0±0.87 |
| PTH (pg.ml<sup>-1</sup>) | 24.6±7.5 | 23.5±7.8 | 25.5(13.6) | 20.5(9.9)** |
| TSH (mIU.l<sup>-1</sup>) | 2.3±1.1 | 1.7±0.8* | 1.2±0.4 | 1.1±0.3* |
| free thyroxine (pmol.l<sup>-1</sup>) | 16.2±0.9 | 16.0±0.1.15 | 18.2±1.7 | 18.8±1.2 |

RMR – resting metabolic rate, RQ – respiratory quotient, CIT – cold-induced thermogenesis (change in RMR), BAT – brown adipose tissue, SUV – <sup>18</sup>F-fluorodeoxyglucose standardized uptake value normalized to lean body mass (LBM), PTH – parathyroid hormone, TSH – thyroid stimulating hormone. Effect of cold exposure was analyzed by two-tailed paired Student's t-test. \*p<0.05, \*\*p<0.01

#### Cohort 3

Here we report on the samples and measurements taken at the baseline visit (before exposure to Fluvastatin) from 11 healthy young men (characteristics - Table 2) participating in the clinical study NCT03189511, more detailed information can be found elsewhere^17^. Maximal induction of NST was aimed to achieve by combining cold exposure with β_3_-adrenergic receptor (AR) agonist (Mirabegron). The volunteers arrived in a fasted state on two separate days with identical schedule to be first screened by indirect calorimetry for an increase in cold-induced thermogenesis of at least 5% of resting energy expenditure. Briefly, volunteers were orally administered 200mg Mirabegron (Betmiga, Astellas Pharma, Switzerland). After 90min rest, standardized cold stimulation with water cooling pads (Hilotherm Clinic; 10°C setting) was commenced lasting another 120min. Afterwards, blood samples were drawn and participants received 75 MBq of ^18^F-FDG intravenously immediately followed PET/MR scan for 50±10 minutes (SIGNA PET/MR, GE Healthcare, Waukesha, WI, USA). Skin and room temperatures were monitored with surface temperature probes. No shivering was reported by any of the participants. At the end of the cooling period, blood samples were drawn from antecubital vein and the participant was transferred to the PET/MR table (SIGNA PET/MR, GE Healthcare, Waukesha, WI, USA). The PET/MR images were acquired from vertex to mid-torso. All scans were performed in the afternoon (between 2-4 pm).

#### Cohort 4

Paired samples of deep neck BAT and adjacent subcutaneous WAT were obtained from patients undergoing neck surgery under general anesthesia, as described in ^18–19^. We analyzed tissues from 36 patients [gender 5M/31F, age 43.5±14.5 years, body mass index 25.9±4.2 kg.m^−2^, waist 87.6±12.8 cm, body fat 32.4±9.1 % (bioelectrical impedance, Omron BF511, Japan), fasting plasma glucose: 4.78±1.00 mmol.l^−1^, triglycerides: 1.35±0.44 mmol.l^−1^, HDL cholesterol: 1.43±0.41 mmol.l^−1^, PTH: 74.15±38.50 pg.ml^−1^]. Anthropometric data are missing from 2 patients. Patients were diagnosed with nodular goiter (n=15), lateral cervical cyst (n=5), hyper functional goiter (n=3), Grave’s disease (n=3), hypothyroidism and nodular goiter (n=2), multi-nodal and diffuse goiter (n=1), hypo functional goiter (n=1), papillary microcarcinoma and Hashimoto’s thyroiditis (n=1), papillary carcinoma (n=1), follicular thyroid adenoma (n=1), parathyroid adenoma (n=1), hyper functional parathyroid adenoma and nodular goiter (n=1), benign vascular tumor (n=1). Patients were on pharmacotherapy with Euthyrox (n=11), L-Thyroxin (n=2), Thyrozol (n=4), Euthyrox + Thyrozol (n=2) or progestin (n=1) and/or with antihypertensives (n=10), vitamin D3/calcium/phosphate supplements (n=6), iron supplements (n=1), hypolipidemics (n=2), insulin pump (type 1 diabetes, n=1). For the analysis of the BAT transcriptome, we only included volunteers with higher relative BAT *UCP1* gene expression than the median *UCP1* expression calculated by combining expression levels in all the corresponding subcutaneous WAT samples (Figure 5A). Plasma samples were collected (n=25) and serum levels of free thyroxine (n=30), free T3 (n=16) and thyroid stimulating hormone (TSH, n=30) were measured.

### Biochemical analysis

To generate serum, collected blood was left at room temperature for 30min, centrifuged (4°C/20min/1200xg) and stored at −80°C. For EDTA/heparin plasma, blood was collected into pre-cooled tubes, centrifuged immediately (4°C/10min/1200xg) and stored at −80°C. Circulating parameters were determined in a certified biochemical laboratory (Alpha Medical, Bratislava, Slovakia) by standardized methods using ADVIACentaur® Immunoassay and ADVIA® Chemistry systems (Siemens, Germany). Reported intact PTH assay specificity: <0.1% cross-reactivity with PTH-related peptide.

### Succinate and lactate analysis

Polar metabolites of plasma were extracted by mixing 20 μl of plasma with 180 μl of 80% methanol. Upon 1 hour incubation at 4 °C, clear extracts were obtained by centrifugation. Non-targeted metabolomics analysis of extracts was performed by flow-injection – time-of-flight mass spectrometry on an Agilent 6550 QTOF system in negative mode ^20^. Metabolite ions were annotated by accurate mass matching using a m/z tolerance of 0.001. Processing and statistical analysis was done in Matlab (The Mathworks, Natick).

### Gene expression

Total RNA was isolated from whole liquid nitrogen-frozen adipose tissue by Trizol extraction, treated by Dnase I and re-precipitated by sodium acetate - ethanol solution, washed in 70% ethanol and dissolved in nuclease-free water for spectrophotometric analysis (Nanodrop 2000). Total RNA was converted to cDNA using High-Capacity cDNA Reverse Transcription Kit or miScript II RT Kit. Fast SYBR™ Green Master Mix and specific primer pairs (5’ – 3’ forward, reverse *UCP1*: TGTGCCCAACTGTGCAATG, GAAGGTACCAACCCCTTGAAGA; *RPL13A1*: GGACCGTGCGAGGTATGCT, ATGCCGTCAAACACCTTGAGA; *PTH1R*: GCCAACTACAGCGAGTGTGTCA, GGTCAAACACCTCCCGTTCA; *PTH2R*: CTTCAACCATAAAGGAGTTGCTTTC, GTGCATAAAATCCCATGTTCCA; *PPARGC1A*: TTACAAGCCAAACCAACAACTTTATC, CACACTTAAGGTGCGTTCAATAGTC; *DIO2*: TCGATGCCTACAAACAGGTG, CATGTGGCTCCCTCAGCTA) designed in Primer Express 3.0 (Thermo Fischer Scientific) were used in quantitative real-time PCR reaction (QuantStudio 5 Real-Time PCR System, Thermo Fischer Scientific). Gene expression was normalized to a housekeeping gene *RPL13A* and to the average relative expression of a gene of interest within all the BAT and WAT samples of a dataset (ddCt). Gene expressions from 4 samples of WAT are unavailable due to being used up for previous experiments (unpaired BAT sample expression values are included in graphs).

### Protein extraction and western blot

Tissue was stored at −80°C and lysate was prepared by homogenizing tissue sample in RIPA buffer (0.5% Sodium Deoxycholate, 2mM EDTA, 150mM NaCl, 1% Triton X100; Sigma-Aldrich) containing protease and phosphatase inhibitors (Sigma-Aldrich) using a mixer mill (Retsch 400MM) and kept at 4°C until lipid and cell debris were removed by centrifugation (15min/12000rpm/4°C). Protein concentrations were measured (BCA assay, Thermo Fisher Scientific). Proteins (30μg) were incubated with loading buffer at 95°C/10min, separated on SDS polyacrylamide gel (12%) and transferred to a PVDF membrane (Merck). After 1 h blocking (Odyssey blocking buffer with 0.1% Tween20), membranes were incubated with primary antibodies against UCP1 (1:500, PA1-24894 Invitrogen), overnight/4°C and against HSP90 beta (1:3000, ab53497 Abcam) and Actin (1:21000, CP01 Calbiochem) for 2h/room temperature. Next, secondary goat anti-rabbit IRDye 680RD or anti-mouse IRDye 800RD antibodies (1:10000, LI-COR) were used to visualize protein of interest (Odyssey Infrared Imaging System, LI-COR). Signal for UCP1 was normalized to the average of the signals for HSP90 and Actin.

### Statistical analysis

All data sets were tested with Shapiro-Wilk normality test. In case of normal distribution (p>0.05), paired Student’s t-test was used and in case of not normal distribution, Wilcoxon matched-pairs signed rank test was used to evaluate the statistical effects of paired samples. Associations between two variables were analyzed by linear regression and Pearson’s correlation coefficient (r) was calculated. Data without normal distribution were logarithmically transformed before correlation analysis. Statistical analyses were performed in GraphPad Prism 6 Software Inc and JMP 4.0.4 (SAS). Data with normal distribution are expressed as mean±SD and data without normal distribution as median (interquartile range - IQR). In cases of wide distribution (TSH, PTH, ddCt UCP1, ddCt PTH2R), graphs were constructed with log Y-scale; negative or zero values are not shown in some graphs (Figure 5A: n=6, Figure 5G: n=2). Outlier values outside the mean±3 x SD range were omitted from analysis in Figure 5F (n=1) and Figure 5G (n=1). Non-physiological values of respiratory quotient (RQ>1) were excluded (Cohort 3, n=1) Cold-induced changes in circulating lactate and succinate were analyzed by Student’s paired t-test and false discovery rate-adjusted p-value was calculated by Benjamini-Hochberg correction.

## Results

### Acute effects of ice-water swimming on levels of parathyroid and thyroid hormones in cold-acclimatized individuals (Cohort 1)

In order to identify changes in hormones and metabolic substrates induced by acute cold exposure in cold-acclimatized volunteers, we analyzed blood samples taken before and within 30 min after completion of ice-water swimming. Ice-water swimming acutely increased serum thyroid stimulating hormone (TSH, Figure 2A) and total thyroxine (T4) was also slightly elevated (Figure 2B), while free T4 (fT4) was decreased (Figure 2C). These data indicate that ice-water swimming is a powerful stimulus affecting the hypothalamic-pituitary-thyroid axis. Furthermore, higher weekly ice-water swimming frequency was associated with a larger cold-swimming-induced decrease in fT4 (Figure 2D), suggesting a faster or more effective fT4 tissue uptake during cold exposure in individuals who are better acclimatized to cold.

**Figure 2:**
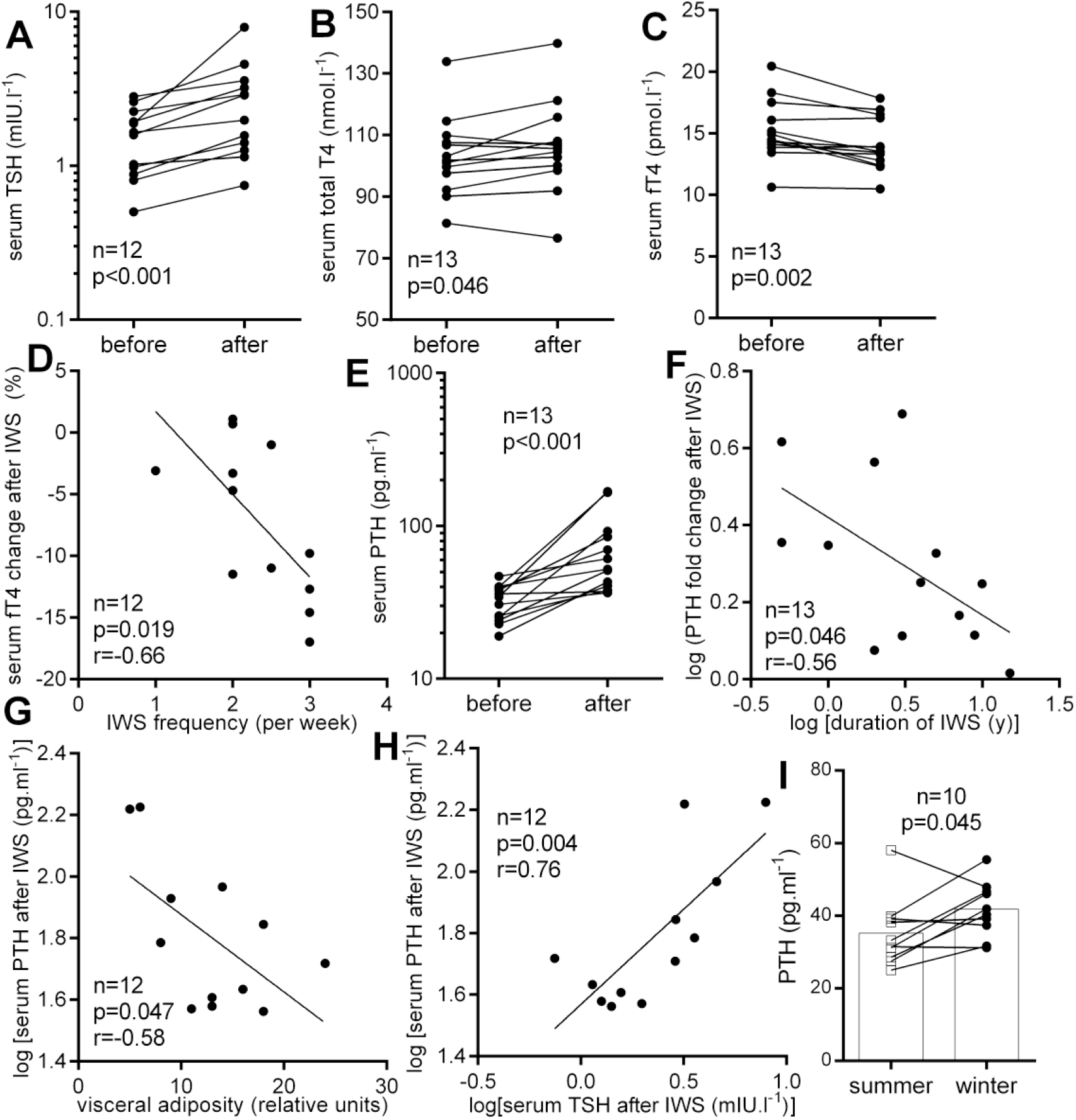
Cold affects circulating parathyroid and thyroid hormones in ice-water swimmers (Cohort 1). Acute effect of ice-water swimming (IWS) on serum (A) thyroid stimulating hormone (TSH), (B) total and (C) free thyroxine (fT4) and (D) its relationship with IWS frequency; acute effect of IWS on (E) parathyroid hormone (PTH) levels and (F) its relationship (F) number of years dedicated to IWS habit, (G) visceral fat and (H) with TSH; (I) PTH seasonal difference. Two-tailed Student’s paired t-test (or Wilcoxon matched-pairs signed rank test and linear regression were used to statistically analyze data. r – Pearson’s correlation coefficient.

Serum PTH was markedly elevated after ice-water swimming, on average by more than 78% (Figure 2E). Furthermore, the cold-water swimming-induced increase in PTH was negatively associated with the number of years dedicated to ice-water swimming habit (Figure 2F), indicating that the response of parathyroid gland to acute cold exposure is less pronounced after several years of cold acclimatization. In addition, PTH levels induced by acute cold exposure were negatively associated with visceral adiposity (Figure 2G) and positively with concomitantly regulated TSH levels (Figure 2H).

### Acute effects of ice-water swimming on insulinemia and circulating metabolites in cold-acclimatized individuals (Cohort 1)

Ice-water swimming acutely increased serum glycemia in 9 from 11 individuals (Figure 3A), while decreasing insulinemia (Figure 3D). Ice-water swimming also induced an acute unanimous increase of lactate (Figure 3B) and succinate (Figure 3C). More importantly, levels of both metabolites correlated positively with PTH (Figures 3E-F).

**Figure 3:**
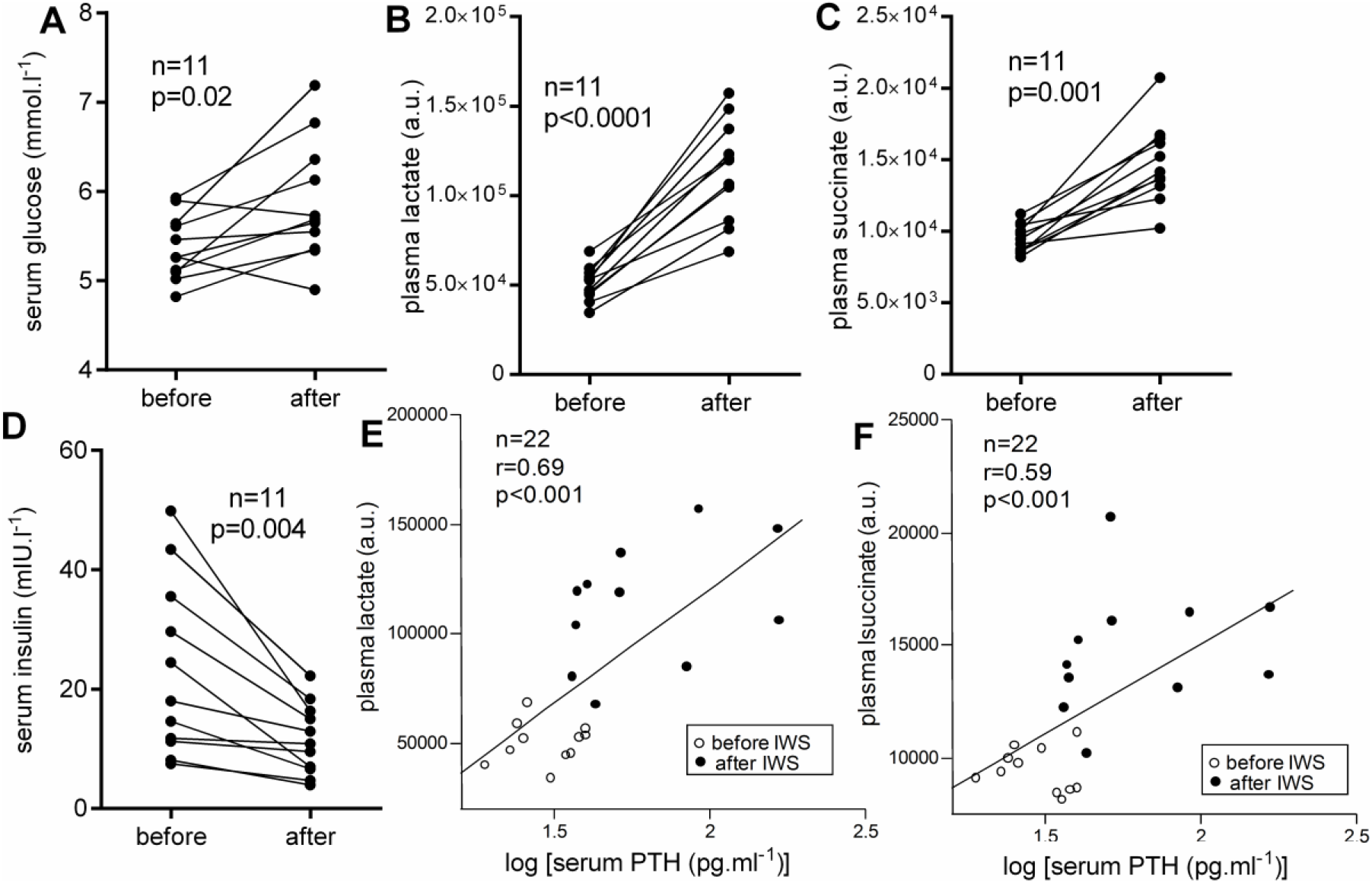
Ice-water swimming modulates substrate metabolites in ice-water swimmers (Cohort 1). Acute effects of ice-water swimming (IWS) on peripheral (A) glycemia, (B) lactate and (C) succinate levels and (D) insulinemia; (E-F) associations of circulating metabolites with parathyroid hormone (PTH) levels during ice-water swimming. Paired Student’s t-test (A, B, D) Wilcoxon matched-pairs signed rank test (C) and linear regression (E-F) were used to analyze data; r – Pearson’s correlation coefficient.

### Seasonal variations in anthropometric, cardiovascular, metabolic and hormonal markers in cold-acclimatized individuals (Cohort 1)

There were no statistically significant differences in cardiovascular and anthropometric parameters of ice-water swimmers obtained in winter and summer season (Table 1). From all the measured parameters, only PTH was significantly higher in winter (Table 1). Figure 2I shows that this was true for majority of examined individuals. It is noteworthy that overweight, obesity and/or increased abdominal adiposity in most of the examined ice-water swimmers were not associated with metabolic disease (Table 1).

### Effects of cold-inducing non-shivering thermogenesis on metabolic preference and circulating levels of thyroid and parathyroid hormones in cold-acclimatized (Cohort 2) and non-acclimatized (Cohort 3) individuals

The induction of non-shivering thermogenesis (NST) was confirmed by increased resting metabolic rate (RMR) in both studied populations (Figure 4A). It is important to note, that while cold-acclimatized ice-water swimmers responded to NST-inducing cold by increasing whole body metabolic preference for lipids (lowering RQ), non-acclimatized men increased metabolic preference for glucose (increased RQ) in response to NST-inducing cold and β_3_AR agonist treatment (Figure 4B). Power of this observation is limited by the fact that acclimatized individuals. Cold exposure aimed at inducing NST decreased TSH in both studied populations (Table 2), and fT4 was not significantly regulated (Table 2). Induction of NST failed to produce any significant PTH response in ice-water swimmers, but the combined cold and β_3_AR-induced stimulation of NST in non-acclimatized individuals decreased circulating PTH levels (Figure 4C).

**Figure 4:**
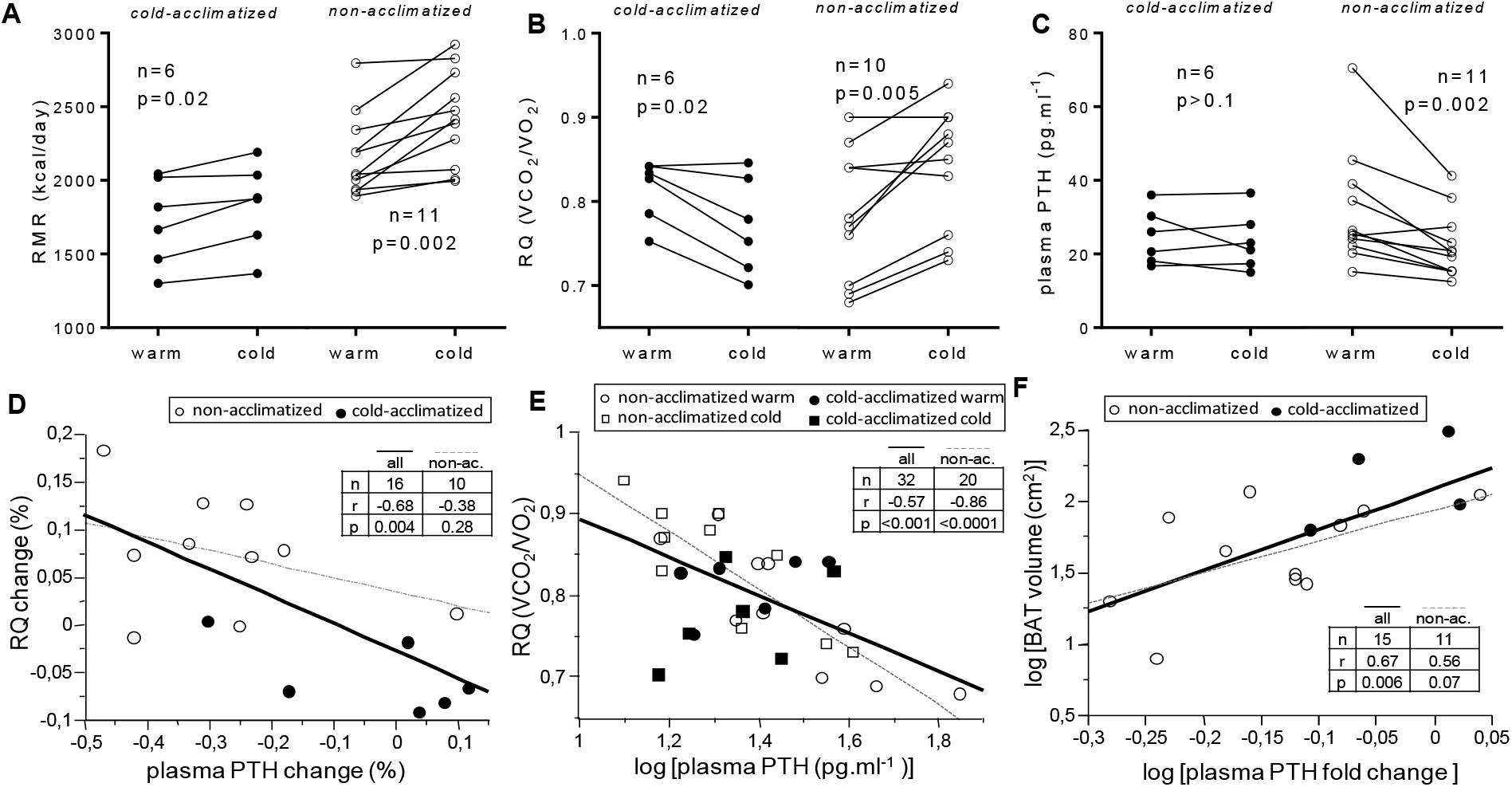
Cold-induced non-shivering thermogenesis distinctly modulates circulating parathyroid hormone in cold-acclimatized and non-acclimatized individuals. Effect of non-shivering cold exposure (Cohort 2) and combined cold/β_3_-adrenergic stimulation (Cohort 3) on (A) the resting metabolic rate (RMR) (B), metabolic substrate preference (respiratory quotient - RQ) and (C) circulating parathyroid hormone (PTH); Relationships between PTH and (D, E) RQ and (F) brown fat volume (the only woman in the group was removed from the correlation). (D-F) fit lines distinguish correlations in all individuals (full line) or within non-acclimatized (non-ac.) group only (grey dashed line). Two-tailed Student’s paired t-test or Wilcoxon matched-pairs signed rank test and linear regression were used to statistically analyze data. r – Pearson’s correlation coefficient.

**Figure 5:**
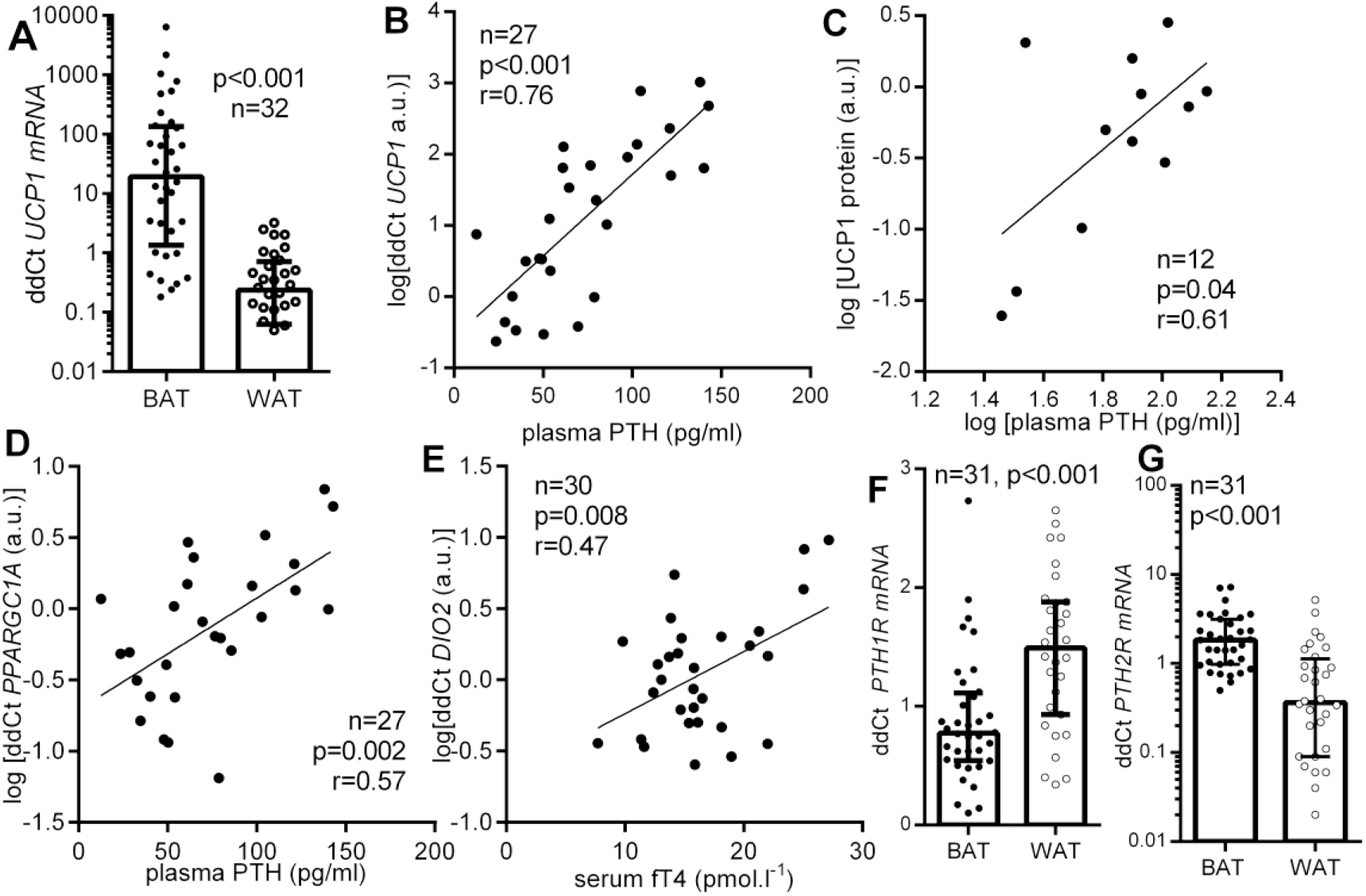
Parathyroid hormone levels are related to human brown fat thermogenic program (Cohort 4). (A) the relative gene expression of uncoupling protein 1 (UCP1) in deep neck brown (BAT) versus subcutaneous white adipose tissue (WAT), (B, C) the relationship between circulating parathyroid hormone (PTH) levels with BAT UCP1 mRNA and protein levels and (D) with the gene expression of peroxisome proliferator-activated receptor gamma coactivator 1-alpha (*PPARGC1A*); (E) association between circulating free thyroxine (fT4) and type 2 deiodinase (*DIO2*) gene expression in BAT; (F, G) gene expression levels of parathyroid hormone receptors (*PTHR*) in BAT and WAT displayed as median with interquartile range, analyzed by Wilcoxon matched pairs signed-rank test.

Next, we explored correlations between circulating PTH, TSH and fT4 and (i) CIT (cold-induced change in RMR), (ii) whole-body metabolic preference for lipids and (iii) BAT glucose uptake and volume measured before and after the cold exposure aimed at inducing NST (cohorts 2 and 3). Most importantly, by including both cold-acclimatized and non-acclimatized individuals we showed that NST-associated change in PTH correlated with the change in whole-body metabolic preference for lipids (Figure 4D). Furthermore, combining levels of PTH detected in warm and cold environment in both studied populations allowed us to evaluate the association between circulating PTH and the whole-body metabolic preference for lipids, confirming presence of the relationship in combined cohorts, as well as in non-acclimatized individuals (Figure 4E). In addition, both baseline (n=11, r=0.71, p=0.01) and cold-regulated levels of TSH (n=11, r=0.82, p=0.002) in non-acclimatized individuals correlated positively with the CIT and cold-induced change in fT4 in non-acclimatized individuals was negatively associated with baseline RMR (n=11, r=−0.73, p=0.01).

### Relationship of PTH and thyroid hormones with thermogenic capacity markers in brown and white fat of non-acclimatized individuals (Cohort 4)

We also explored the links between PTH and brown adipocyte gene/protein markers in deep-neck BAT and adjacent subcutaneous WAT from non-acclimatized patients undergoing elective neck surgery, who were not acutely challenged with cold (Cohort 4). Expression of *UCP1* mRNA in BAT showed large variability (Figure 5A) but observed variability was paralleled by corresponding variability in PTH. Most importantly, fasting plasma PTH levels positively correlated with deep-neck BAT gene expression of *UCP1* (Figure 5B)*, PPARGC1A* (Figure 5D) and *DIO2* (n=27, p=0.03, r=0.43) as well as with UCP1 protein levels in BAT (Figure 5C). The importance of paracrine thyroid activity on adipose tissue was indicated by positive association between deep-neck BAT expression of *DIO2* and circulating levels of fT4 (Figure 5E).

Furthermore, we found that both PTH receptors (*PTH1R, PTH2R*) were expressed in brown and white fat (Cohort 4). While *PTH1R* gene expression was significantly higher in WAT, *PTH2R* showed higher expression in BAT (Figure 5F-G). Expression levels of *PTH1R* paralleled those of *PTH2R* in BAT (n=36, p=0.03, r=0.37), but the relationship was not present in WAT, indicating depot-specific differences in PTH signaling in humans.

## Discussion

The mechanisms of cold-induced metabolic and thermogenic activation of BAT have been extensively studied in recent years due to the promising therapeutic potential in the prevention and treatment of obesity-related metabolic diseases. In our study, we have observed distinct cold-induced changes in circulating humoral factors, which are potentially implicated in the regulation of adaptive thermogenesis.

This work clearly shows that while cold-stress-associated ice-water swimming induces profound increase in circulating PTH in cold acclimatized individuals, it remains unchanged when milder cold stimulus aimed at inducing NST is applied to similarly acclimatized individuals and that PTH decreases under similar NST-inducing conditions in individuals not acclimatized to cold. To our best knowledge, this is the first evidence showing acute cold exposure-related changes of PTH. Our data suggest that PTH response might be modulated by both the intensity of cold stimulus, the level of cold acclimatization as well as by the between the magnitude of ice-water-induced change in PTH and fT4 and the duration of this habit and frequency of ice-water swimming sessions, respectively. It could be speculated that distinct and perhaps more efficient adaptive thermogenic mechanisms develop in experienced (better acclimatized) ice-water swimmers. This notion is supported by the evidence of altered thermoregulatory responses present in ice-water swimmers during cold water immersion compared to controls ^21^. We observed that cold-induced thermogenesis seemed to be lower in cold-acclimatized individuals, which is in line with the previous reports^21^, however BAT volume defined by the ^18^FDG uptake seemed to be similar. We also observed that whole-body metabolic preference for lipids increases upon NST-stimulating cold exposure in ice-water swimmers while non-acclimatized individuals increase their preference for glucose under similar conditions. This is in line with the observation that non-shivering thermogenesis is in young healthy BAT positive individuals associated with similar increase of respiratory quotient^3^. This may suggest that regular ice-water swimming is associated with an adaptive change in the cold-induced whole-body metabolic preference for lipids.

The differential regulation of circulating PTH in cold-acclimatized individuals subjected to ice-water swimming and to non-shivering cold exposure could stem from the intensity of the cold stimulus, from the exercise component of ice-water swimming ^22^, from the post-swimming shivering thermogenesis and from the difference in the body area exposed to cold (full-body cold water immersion *vs.* cooling pads). Meanwhile, the difference in cold-induced PTH regulation between cold acclimatized and non-acclimatized individuals is most likely related to the adaptive response to repeated cold water immersion, although we cannot rule out the effect of β_3_AR agonist and the fact that the shivering threshold in cold acclimatized individuals lies at much lower core body temperature ^21^.

Present evidence indicates that PTH or PTH-related protein (PTHrP) drive the thermogenic program in primary mouse brown, beige and white adipocyte cultures, as well as in brown, inguinal and epididymal white adipose tissue depots of mice ^16, 23–24^. This raises the question whether PTH could be involved in regulation of adaptive thermogenic process in humans. A recent study has shown the browning, thermogenic and lipolytic effects of PTH in primary cultures of human subcutaneous white adipocytes ^15^. Here, we provide several pieces of evidence that PTH might specifically modulate metabolic response to cold stress and/or BAT thermogenic capacity in cold acclimatized humans. We found that the cold-induced metabolic preference for lipids, which specifically increased in cold-acclimatized individuals, was associated with the acute change in PTH. Therefore, we speculate that the ice-water swimming-induced increase in circulating PTH could potentially be linked with the induction of whole-body fat utilization. It is important to note that the baseline levels of PTH in cold-acclimatized ice-water swimmers were more than 18% higher in the winter than in the summer season. This could reflect presence of adaptive changes that might predispose the individuals to more effective cold response during the cold water swimming. We also found a negative association between the ice-water swimming-induced increase of PTH and visceral adiposity, which might be related to the potential of repeated short-term cold-water-induced PTH spikes to increase the adipose tissue lipid utilization. Furthermore, the magnitude of the PTH change associated with the cold-induced NST correlated positively with the BAT volume and PTH levels before and after ice-water swimming strongly correlated with circulating lactate and succinate, the key metabolite markers of BAT thermogenesis ^25–27^. Interestingly, acidity associated with systemic lactate accumulation might trigger the release of PTH ^28^. Unlike in ice-water swimming, induction of NST is not associated with systemic increase of lactate ^25^. Therefore, it is plausible to think that lactate release could be an important factor regulating PTH response to cold although further experiments need to validate this notion. Lastly, systemic levels of PTH in non-acclimatized individuals who were not exposed to cold correlated well with BAT markers *UCP1*, *PGC1α* and *DIO2* mRNA, as well as with UCP1 protein. We propose that this might reflect the natural inter-individual variability in the thermogenic capacity within the group of patients subjected to elective neck surgery. Furthermore, primary hyperparathyroidism in humans is associated with higher prevalence and magnitude of ^18^F-FDG uptake into BAT ^24^ and with elevated expression of several thermogenic genes in deep neck BAT, but not in subcutaneous WAT ^16, 23^. In fact, we showed that human deep neck BAT and subcutaneous WAT express both types of PTH receptors, indicating that PTH might in fact directly regulate adipose tissue functional state. Moreover, PTH1R, a receptor shared by PTH and PTHrP, had significantly higher expression in WAT, while PTH2R, receptor specific for PTH was enriched in BAT. This observation replicated our RNAseq data^19^. Collectively, we believe that these are important findings supporting the physiological role of PTH in cold defense mechanisms, potentially including the adipose tissue metabolic and/or thermogenic activity.

The elevated TSH levels after ice-water swimming and the opposite regulation in response to cold-induced NST in both non-acclimatized and cold-acclimatized individuals is in line with the evidence that severe cold (cold water immersion) has the capacity to increase but mild cold (cold air/cooling pads) results in decreased or unchanged circulating TSH ^29–33^. Similarly, the decrease in fT4 associated with ice-water swimming, which was not found during cold-induced NST, indicates the sensitivity of thyroid axis to cold exposure intensity or duration. This is supported by the observation that *DIO2* expression and activity increases in BAT of healthy men after prolonged mild cold exposure ^4^ that could promote T4-T3 conversion and T4 tissue uptake. In the group of patients with varying thyroid function (Cohort 4), fasted fT4 levels positively correlated with brown adipose tissue *DIO2* mRNA, supporting the sensitivity of human BAT to peripheral thyroid hormones. Furthermore, individual variability in NST-induced change in fT4 was negatively correlated with energy expenditure in non-acclimatized men. Our data provide important new evidence on the regulation of thyroid hormones by cold in acclimatized and non-acclimatized humans and on their role in BAT thermogenesis in humans.

Limitations of the study include the imbalanced gender ratio and age variability between the study cohorts, which reflects the limited capacity to recruit such specific groups (ice-water swimmers, patients undergoing neck surgery). The smaller number of participants in Cohort 2 and several missing values for plasma parameters of Cohort 4 (blood was not collected at the pilot stage of the study) certainly limited the statistical power, but in our honest opinion, presented data provide important evidence on the potential role for PTH in modulating metabolic and thermogenic response to cold in cold-acclimatized individuals. It is important to note that the sequence/timing of the post-swimming blood collection was not associated with any variability in either absolute values or cold-induced changes of PTH, TSH or T4. We also acknowledge some differences in the NST-inducing protocols (use of β_3_-AR agonist, duration of cold and necessity to lower cooling temperature in cold acclimatized individuals) between cohorts 2 and 3 that might reduce the ability to compare the effects between the groups. Complexity of the experimental approach could have been extended by comparing effects of swimming or comparable physical activity without concomitant cold exposure to control for exercise-specific changes, as well as by extending the protocol to study time-dependent dynamics of the cold-induced changes in PTH. However, we believe our study provides important novel evidence creating the grounds for future experiments exploring the role of PTH in human metabolic & thermogenic response to cold.

## Conclusion

We report that circulating parathyroid hormone and thyroid axis components were acutely stimulated by ice-cold water swimming, but not by mild cold exposure inducing non-shivering thermogenesis in cold acclimatized individuals. The relationships between PTH and cold-induced metabolic substrate preference for lipids and the presence of systemic and brown adipose tissue markers of thermogenic process provide pilot evidence indicating that PTH is involved in metabolic and thermogenic response to cold stress in humans.

## Acknowledgments

We would like to thank Prof. Dr. Nicola Zamboni for the metabolomic measurements and all the study volunteers for participating in this study.

## Data Availability

The datasets generated during and/or analyzed during the current study are not publicly available but are available from the corresponding author on reasonable request.

